# Frontoparietal Hubs Leverage Probabilistic Representations and Integrated Uncertainty to Guide Cognitive Flexibility

**DOI:** 10.1101/2025.03.21.644222

**Authors:** Stephanie C. Leach, Hannah Hollow, Jiefeng Jiang, Kai Hwang

**Author notes:** These authors contributed equally to this work (Jiefeng Jiang, Kai Hwang).

## Abstract

Adaptive behavior requires integrating information from multiple sources. These sources can originate from distinct channels, such as internally maintained latent cognitive representations or externally presented sensory cues. Because these signals are often stochastic and carry inherent uncertainty, integration is challenging. However, the neural and computational mechanisms that support the integration of such stochastic information remain unknown. We introduce a computational neuroimaging framework to elucidate how brain systems integrate internally maintained and externally cued stochastic information to guide behavior. Neuroimaging data were collected from healthy adult human participants (both male and female). Our computational model estimates trial-by-trial beliefs about internally maintained latent states and externally presented perceptual cues, then integrates them into a unified joint probability distribution. The entropy of this joint distribution quantifies overall uncertainty, which enables continuous tracking of probabilistic task beliefs, prediction errors, and updating dynamics. Results showed that latent state beliefs are encoded in distinct regions from perceptual beliefs. Latent-state beliefs were encoded in the anterior middle frontal gyrus, mediodorsal thalamus, and inferior parietal lobule, whereas perceptual beliefs were encoded in spatially distinct regions including lateral temporo-occipital areas, intraparietal sulcus, and precentral sulcus. The integrated joint probability and its entropy converged in frontoparietal hub areas, notably middle frontal gyrus and intraparietal sulcus. These findings suggest that frontoparietal hubs read out and resolve distributed uncertainty to flexibly guide behavior, revealing how frontoparietal systems implement cognitive integration.

**Significance:** Flexible human behavior often depends on integrating information from multiple sources, such as memory and perception, each of which can be corrupted by noise. For example, a driver must integrate traffic signals (external cues) with their destination plan (internal goals) to decide when to turn. This study reveals how the human brain integrates multiple information sources to guide flexible behavior. More specifically, distinct brain regions encode internal beliefs and external sensory representations, while frontoparietal regions integrate this information in response to input noise. These findings provide a complete account of how the brain encodes and integrates multiple inputs to guide adaptive behavior.

## Introduction

The ability to engage in complex goal-directed behavior is a hallmark of human intelligence. To achieve goals, humans construct rich cognitive representations that capture rule structures and contextual contingencies (Cellier, Petersen, & Hwang, 2022; Rangel, Hazeltine, & Wessel, 2023). These cognitive representations transform sensory and mnemonic inputs into actions and decisions to guide goal-directed behavior. Critically, this often involves integrating multiple sources of information, including perceptual inputs and internal states. For instance, when driving, flashing lights from a distant accident must be combined with internal beliefs about traffic to select an optimal alternative route. However, the neural and cognitive processes underlying cognitive integration, for combining internal/external inputs, are not well understood. Cognitive integration is challenging because some sources are non-observable (e.g., internal thoughts), corrupted by noise, or stochastic in nature. Thus, the mechanisms for integrating stochastic information from observable and non-observable sources are unknown—leaving a critical gap in our understanding of cognitive integration.

Integrative functions are believed to be supported by hub regions in the frontoparietal network (FPN), which exhibit extensive connectivity across multiple systems (Gratton, Sun, & Petersen, 2018; Power, Schlaggar, Lessov-Schlaggar, & Petersen, 2013; van den Heuvel & Sporns, 2013). Hubs have been proposed to leverage this connectivity architecture to flexibly receive inputs and transmit outputs among diverse brain networks to support integration (Sporns & Betzel, 2016). Studies of functional brain networks often identify frontoparietal regions exhibiting strong hub properties (Bertolero, Yeo, & D’Esposito, 2015). However, connectivity patterns alone do not reveal the underlying neurocognitive mechanisms of integration. Specifically, they neither explicitly test the computational processes enabling integration nor do they clarify the representational structure of the encoded cognitive information. Consequently, how hub regions integrate diverse information remains unclear.

To address this, we propose a computational framework clarifying the integration of internally maintained or externally cued stochastic information. This framework specifies: (1) inputs-to-be-integrated, (2) processes transforming inputs into an integrated representation, (3) outputs derived from this representation, and (4) updating mechanisms modifying inputs based on the integrated outputs. Critically, incorporating these elements with neuroimaging can precisely probe underlying neural mechanisms.

We developed a Bayesian-based computational model to infer latent variables that represent subjective beliefs from external and internal information sources (inputs), as well as from the output of their integration. This model posits that internal and external sources of information are integrated into a joint probabilistic distribution that maps onto a conditional probabilistic distribution (Fig. 1). The overall uncertainty in integration is measured through the entropy of the joint distribution (Fig. 1A-C). The probability of performing a given action/decision is conditioned on the joint contribution of both external and internal sources (Fig 1D). The latent variables include both the categorical beliefs of inputs/outputs and their associated uncertainty, which enables further analysis of prediction errors and updating processes.

**Fig 1.**
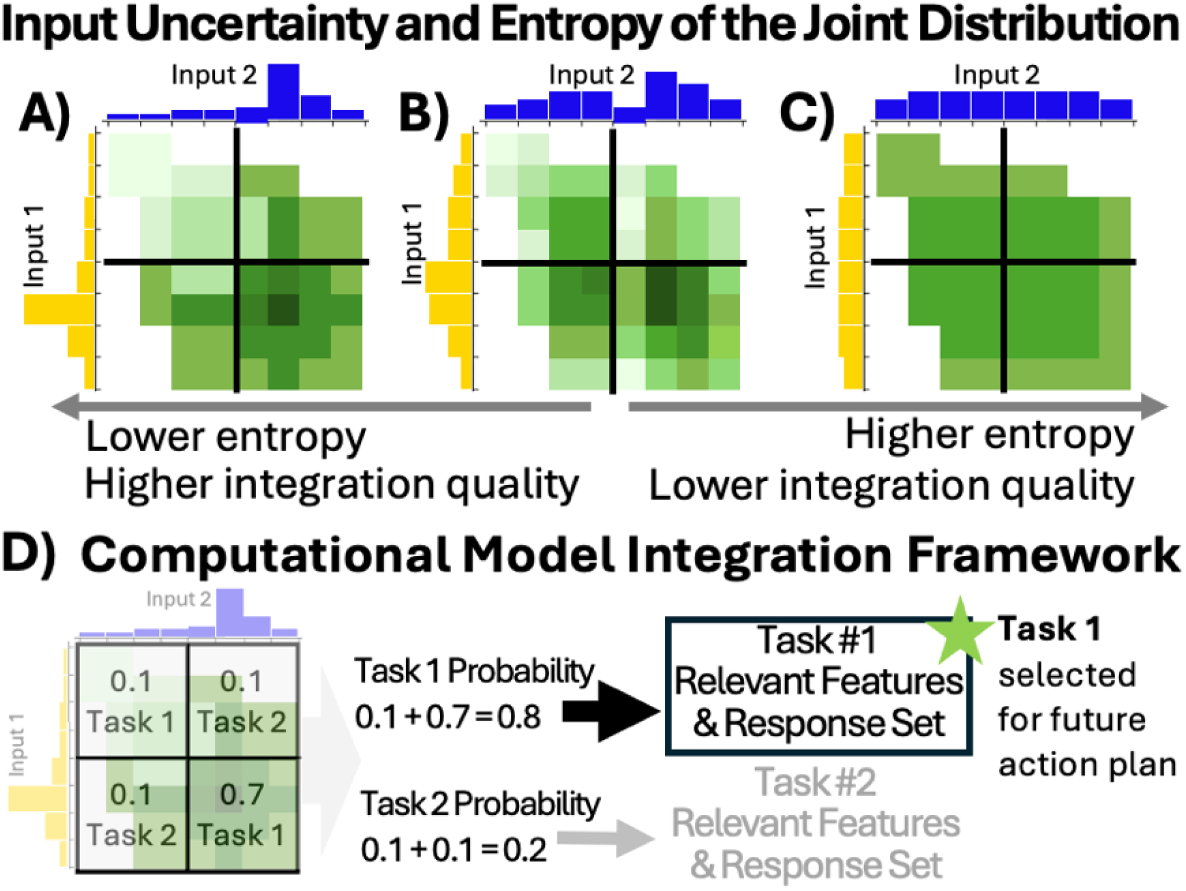
Integrated Uncertainty. (**A-C**) Depiction of how integrated uncertainty, or entropy of the joint distribution, fluctuates with different levels of input uncertainty. The shading of the cells in the joint distribution reflects the peak of the integrated distribution. A clear peak, as indicated by a darker shade of green, indicates a low level of integration uncertainty. A lack of a clear peak indicates a high level of integration uncertainty. (**A**) Low levels of input uncertainty lead to higher integration quality and therefore lower levels of integrated uncertainty. (**B**) Moderate levels of input uncertainty lead to moderate levels of integrated uncertainty. (**C**) High levels of input uncertainty lead to a lower integration quality and therefore high levels of integrated uncertainty. (**D**) Depiction of integration framework. Here, two tasks are associated with a joint distribution of two stochastically varying input sources of information. This 2×2 joint distribution maps onto a conditional probabilistic distribution to predict a should be executed task (Task 1 vs. Task 2). Every cell in this 2×2 grid shows the likelihood that the current inputs fall within each cell’s defined range. To decide which task to execute (Task 1 or Task 2), the model sums probabilities of the cells associated with each task and chooses the task whose cells sum to the higher value.

Our framework provides trial-by-trial predictors detailing input-integration-output representational structures, uncertainty estimates, and prediction errors driving belief updating. We combined model-derived estimates with fMRI analyses to comprehensively investigate the neurocognitive processes underlying the integration of stochastic information. This allowed us to investigate both the processes enabling integration and the representational structure of encoded cognitive information. Results show latent state beliefs are encoded primarily in middle frontal gyrus (MFG), mediodorsal thalamus, and inferior parietal lobule (IPL), whereas perceptual beliefs are encoded primarily in lateral temporo-occipital regions, intraparietal sulcus (IPS), MFG, and superior frontal gyrus (SFG). The integrated joint probability of the external/internal input beliefs and their integrated uncertainty (entropy) converged in frontoparietal hubs. In summary, distinct information sources are represented in largely separate brain systems but converge in frontoparietal hubs, where entropy provides a mechanism for resolving distributed uncertainty and guiding adaptive cognitive control.

## Materials and methods

### 2.1 Experimental design

To address cognitive integration, we utilized an experimental design in which participants integrated one external and one internal source of information to determine the correct task for each trial (Fig. **2**). We developed a Bayesian model (Fig. **3A**) to capture computational processes underlying cognitive integration in this experimental design. Participants completed an fMRI scanning session while performing this cognitive integration task. Before scanning, all participants completed a tutorial and practice session. Scanning began after participants achieved a minimum practice score of 80%. We collected five functional runs, each containing 40 task trials, resulting in a total of 200 trials across the session. Each functional run lasted approximately 8.5 minutes.

**Fig 2.**
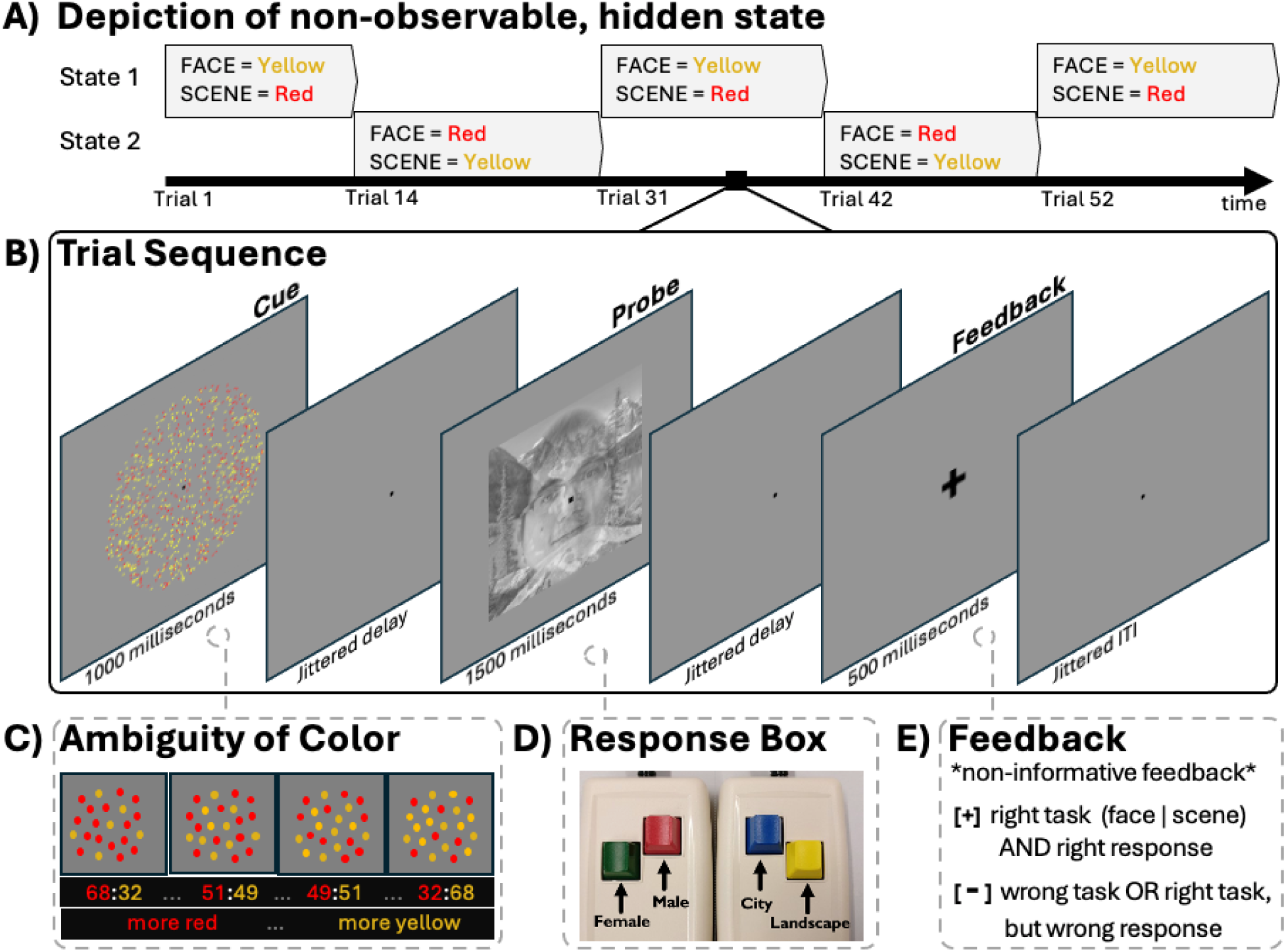
Experimental Design. (**A)** Depiction of an example non-observable, volatile state transition that changes every 10-20 trials. **(B)** Trial sequence: cue screen (1000ms), jittered delay (range: 0.5-10.5 seconds, average=3.75 seconds), probe screen (1500ms), jittered delay (range: 0.5-10.5 seconds, average=3.75 seconds), feedback screen (500ms), and jittered intertrial interval (range: 0.5-10.5 seconds, average=3.75 seconds). (**C)** Depiction of color ambiguity manipulation. Some trials were less ambiguous with proportions closer to 68:32 while other trials were more ambiguous with proportions closer to 51:49. **(D)** Response setup for fMRI. Note, responses for each task were split by hand. Face task (left hand) discrimination judgement response options were female (middle finger) or male (index finger). Scene task (right hand) discrimination judgement response options were city (index finger) or landscape (middle finger). (**E**) Possible feedback. Note, feedback was non-informative—it did not clarify if participants were wrong because they picked the wrong response within the correct task or because they did the wrong task. Example face stimulus is a photo of an author of this manuscript.

**Fig 3.**
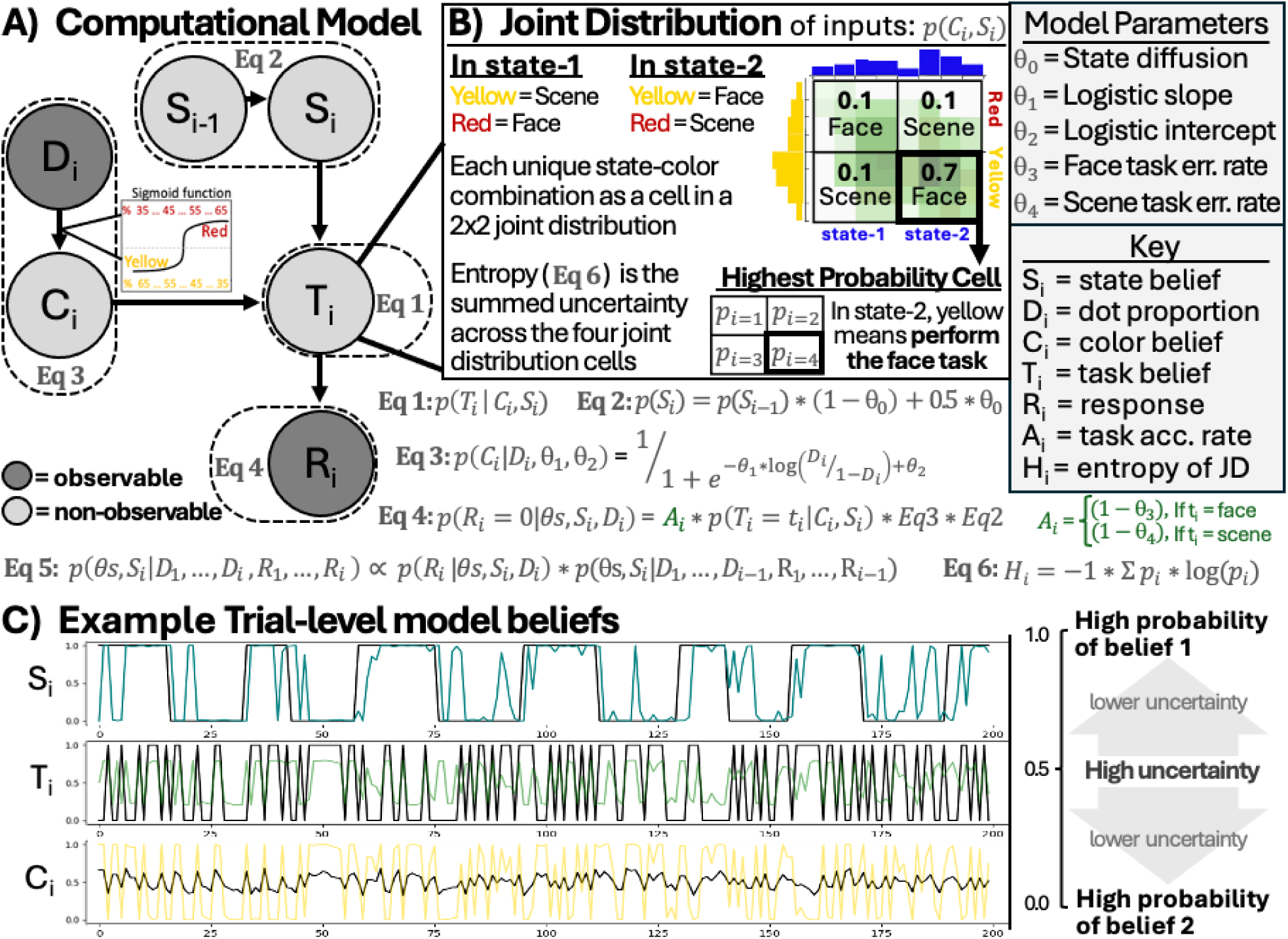
Computational Model. (**A**) Computational (Bayesian generative) model. Observable information (R_i_ and D_i_) was used to infer three latent variables: state belief (Si), color belief (Ci), and task belief (Ti). A sigmoid function (see Equation 3) was applied to the presented color proportion (D_i_) to get an estimate of color belief (C_i_). The state belief S_i_ was inferred from D_i,_ and R_i_. To allow the previous state to influence the current state, trial-wise diffusion of S_i_ was implemented (see Equation 2). S_i_ and C_i_ were integrated into a 2×2 joint distribution from which the task belief T_i_ was inferred (see Equation 1). (**B**) In the example of a trial-level 2×2 joint distribution shown here, the probability that the face task was the task-to-perform is 0.8, summed from 0.1 (state = 1 and color = red) + 0.7 (state = 2 and color = yellow). Gray boxes on the far right side indicate the free model parameters fit during the inference procedure (see Equations 4-5) and a legend clarifying letter codes used in the model depiction and equations. These parameters include a diffusion parameter (allowing S_i–1_ to influence S_i_), a logistic function intercept and slope (capturing color bias and the high-uncertainty range, respectively), face and scene task error rates (used for updating following incorrect feedback; see Methods section 2.4 for full details). (**C**) Trial-level model estimated beliefs from one example subject. Black lines: true state, task, or color. Blue, green, and yellow lines: state belief, task belief, and color belief, respectively. The closer belief values are to 1 or 0, the more certain the belief, while the closer it is to 0.5, the more uncertain the belief.

### 2.2 Participants

We recruited 49 healthy participants (19 male and 30 female, *M*_age_ = 22.396, *SD*_age_ = 4.372, *Range*_age_ = 18-35 years) from the University of Iowa and surrounding area to participate in this study. All participants had normal or corrected to normal visual acuity and color vision, were right-handed, and reported no history of epilepsy, psychiatric, or neurological conditions. This study was approved by the University of Iowa Institutional Review Board and conducted in accordance with the principles outlined in the Declaration of Helsinki. All participants gave written informed consent. Out of these 49 participants, 11 were excluded for excessive movement and/or chance-level behavioral performance during fMRI scanning, resulting in a final sample of 38 participants (15 male and 23 female, *M*_age_ = 22.95, *SD*_age_ = 5.88, *Range*_age_ = 18-35 years).

### 2.3 Experimental tasks

Each trial (Fig. **2B**) began with the presentation of an array of red and yellow dots moving randomly in a circle (cue screen; presented for 1000-milliseconds). During this presentation, participants determined if there were more red or yellow dots in this array. The proportion of dots for the dominant color was drawn from a continuous range of .51 to .68. After a jittered delay (average duration of 3.75 seconds; range of 0.5 – 10.5 seconds), a probe screen appeared for 1500 milliseconds, showing a partially transparent face overlaid on a partially transparent scene image. At this point participants performed either the face task (report whether the face was male or female; Fig. **2D**) or the scene task (report whether the scene depicted a city or a nature landscape; Fig. **2D**). Response mappings were assigned to separate hands: the left hand for the face task (middle finger for female, index finger for male) and the right hand for the scene task (index finger for cities, middle finger for nature scenes). These mappings were fixed and not counterbalanced across subjects. As previously described, participants needed to integrate the hidden state (Fig. 2A) and perceptual color information (Fig. 2C) to determine which task to perform (Fig. 2D). In one hidden state, more red dots indicated that the correct task was the face task, while more yellow dots indicated the correct task was the scene task. In the other hidden state, this pairing was reversed: more red dots indicated the scene task, and more yellow dots indicated the face task. For every participant, the hidden state covertly switched every 10–20 trials, resulting in 12 switches over the course of the experiment (identical for all participants). After a second jittered delay (average duration of 3.75 seconds; range of 0.5 – 10.5 seconds), a feedback screen appeared for 500 milliseconds, showing either a plus or minus sign. A plus sign indicated that the participant performed the correct task and made the correct discrimination judgement (male/female or city/landscape), while a minus sign indicated either the participant performed the incorrect task and/or made the wrong discrimination judgement (Fig. 2E). Finally, each trial ended with a jittered inter-trial-interval (average duration of 3.75 seconds; range of 0.5 – 10.5 seconds).

### 2.4 Computational model

Here, we outline a Bayesian model (Fig. **3A**) that captures the computational processes underlying cognitive integration in this paradigm. This model estimates two probabilistic distributions, a hidden state and a perceptual belief, which are integrated into a joint probability distribution. From this joint distribution, the model generates a probabilistic task-output belief specifying which task the participant should perform. In other words, this model generates trial-by-trial predictors (Fig. **3B**) for input representations, their integrated product (i.e., joint probability distribution), the integrated uncertainty (entropy), and the resulting task output representation. These trial-by-trial predictors can then be combined with neuroimaging analyses (sections 2.8.2–2.8.4) to probe the neural underpinnings of the integration of probabilistic, observable and non-observable sources of information.

In the Bayesian model (Fig. 3A), for trial i, the probabilistic state belief (S_i_, representing the two color-task mappings in Fig. 2A) and probabilistic color belief (C_i_) are integrated into a 2×2 joint distribution, which was used to infer the probabilistic task belief (T_i_, with Task 1 indicating the face task and Task 2 indicating the scene task).

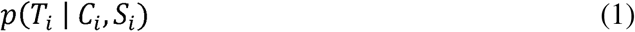

Here, state belief (S_i_), color belief (C_i_), and task belief (T_i_) are latent variables and state belief (S_i_) and color belief (C_i_) are inferred from the directly observable color proportion (D_i_) and response made by the participant (R_i_). Because the previous state belief (S_i-1_) should influence the current state belief (S_i_), which should in turn influence the subsequent state belief (S_i+1_), a trial-wise diffusion of S_i_ was implemented with the following equation. Note, this diffusion is done at the start of each trial (i) and so S_i_ is on both the left and right side of the updating rule.

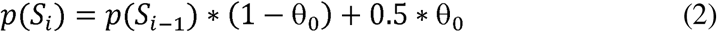

In other words, at the beginning of each trial, a confusion in the form of random guess (i.e., 0.5 probability) of the state is introduced to the previous state belief with a weight of θ_0_. By applying this theta weight to the state belief, the state belief is brought closer to 0.5. How much closer to 0.5 the state belief gets depends on the specific value of θ_0_. The range of possible change in state belief ranges from 0 (i.e., no confusion) to 0.5 (high confusion) (Behrens, Woolrich, Walton, & Rushworth, 2007; Jiang, Heller, & Egner, 2014).

Color belief (C_i_) was estimated from the observable color proportion (D_i_) using a sigmoid function, see equation 3.

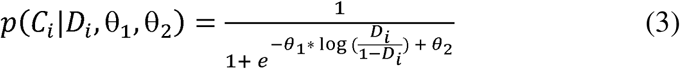

The two free parameters (θs) estimating this sigmoid function determined the (1) slope (accounting for color uncertainty/perception) and (2) intercept (accounting for color bias).

The model predicts the response by combining the term p(T_i_ | C_i_, S_i_) with two additional free parameters, the participant’s respective error rates for the face (θ_3_) and the scene tasks (θ_4_), to predict the responses (R_i_). R_i_ was coded as 0 for correct task and correct response or 1, which could mean either correct task but incorrect response or incorrect task and either correct or incorrect response. Taken the above steps together, for the true task t_i_, the prediction of generating the correct response given the free parameters, the belief of state and color input [i.e., *p*(*R_i_* = 0|*θ_S_, S_i_, D_i_*)], is estimated in the following manner:

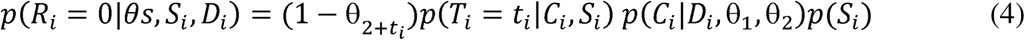

Where 2+t_i_ indicates the error rate corresponding to the true task (either face or scene task depending on the current true task). The prediction of committing an error [i.e., *p*(*R_i_* = 1|*θ_S_, S_i_, D_i_*)] can be obtained using 1 − *p*(*R_i_* = 0|*θ_S_, S_i_, D_i_*). This step of predicting R_i_ despite R_i_ being an observable variable was necessary to estimate the participant’s respective error rates for the face (θ_3_) and the scene (θ_4_) tasks, which are necessary for trial-to-trial belief updating.

By using the trial-wise values of the directly observable variables (D_i_ and R_i_), the generative model was inverted to infer latent variable distributions (Fig. 3A).

For the first trial, the prior is initialized as a uniform distribution. As the trials progress, the posterior of the previous trial becomes the prior for the current trial. Updating the prior was achieved by multiplying the likelihood (*p*(*R_i_*|*θ_S_, S_i_, D_i_*)) and prior (i.e., the posterior of the previous trial, *p*(*θ_S_, S_i_|D*_1_, …, *D_i_*_-1_, *R*_1_, …, *R_i_*_-1_)) and dividing them by a normalization constant (Equation 5).

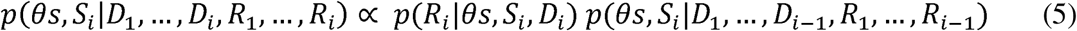

As shown in Equation 5, the inference procedure estimates the probability distribution for each free parameter. In the implementation of the model, each probability distribution is coded as a vector representing the probability at different values. We tested the logistic function’s slope parameter over a range of 1.5–5, based on model fits to independently collected pilot behavioral data (not reported here). The range of values tested for the logistic function intercept parameter was log(0.45/0.55) to log(0.55/0.45), corresponding to actual color biases up to a 55:45 (dominant:non-dominant color) split. Error-rate parameters for both the face and scene tasks were tested between 0 to 0.35, based on potential error rate ranges for each task. Finally, the range tested for the parameter associated with the diffusion of state information across trials was 0 to 0.9. This range was also selected based on model fits in a pilot sample not reported here. We then used the softmax function to select the optimal value for each of the five theta parameters for each participant.

The proposed model treats both state belief (S_i_) and color belief (C_i_) as probabilistic representations (Fig. 3B) for. As a result, the 2×2 joint probability distribution encoded the combined uncertainty of these two sources of inputs. Moreover, because the joint probability distribution encodes the combined uncertainty of the inputs, the output, Ti, that is generated from the joint probability distribution is also probabilistic (Fig. 3B). State belief must be maintained across trials and updated with feedback. Equations 4 and 5 specify this update, including theta parameters for the respective error rates for the face (θ_3_) and the scene tasks (θ_4_).

#### 2.4.1 Uncertainty Measures for Model Beliefs

The proposed model infers three key latent variables representing information participants used to accomplish integration, two representing the inputs (state and color) that are integrated and one representing the output of integration (task). Critically, these three latent variables are probabilistic and therefore contain not only the categorical belief (e.g., red or yellow), but also the uncertainty associated with this belief on each trial. Uncertainty estimates can therefore be extracted from these latent variables to more fully capture the representational structure (see paragraph below for more details on how uncertainty was extracted). Moreover, the 2×2 joint probability distribution encodes not only the specific categorical combinations of the input (state and color) beliefs, but also the overall uncertainty of the join distribution (entropy), which we also used in subsequent neuroimaging analyses.

Latent variable uncertainty estimates for subsequent neuroimaging analyses included both the uncertainty of inputs for integration (state and color beliefs) and output (task belief), as well as the uncertainty associated with the updating mechanisms (prediction errors and derivatives). We extracted uncertainty from each of the three latent variables (state belief, S_i_; color belief, C_i_; task belief, T_i_) using the following steps. First, each belief was centered by subtracting 0.5 from the values. Second, the absolute value of the centered belief was taken to represent the magnitude of uncertainty. Finally, for ease of interpretation, the signs of these uncertainty scores were inverted. This inversion aligned the uncertainty scores for the three latent variables with the overall uncertainty of the joint distribution (entropy), such that values closer to zero indicated lower, and higher values indicated greater uncertainty.

Prediction errors were calculated by taking the absolute value of the difference between each model’s estimated belief and its true value. Here, lower values indicate that the participant’s belief closely matched the true value, while higher values reveal substantial mismatches between the participant’s belief and the true value. Calculating prediction errors this way provided a continuous measure of prediction error where the magnitude reflected how much the participant’s belief deviated from the true value. Finally, state derivative, which assessed the magnitude of the change in state belief from trial to trial, was calculated by subtracting the state belief on trial N-1 from trial N and taking the absolute value. This captured how much the state belief shifted from one trial to the next, with larger values reflecting greater belief revision.

Finally, the overall uncertainty of the joint distribution was calculated using Shannon’s entropy (Equation 6):

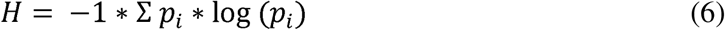

Here, p_i_ refers to each cell in the 2×2 joint distribution. Shannon’s entropy was used to quantify the overall uncertainty across the four possible outcomes in this joint distribution.

### 2.5 Computational model comparisons

This computational model assumes probabilistic representations for both state belief (S_i_) and color belief (C_i_), which are integrated into a joint distribution to determine the task belief (T_i_). To test its validity, we compared this model against three alternative models: (1) a model with dichotomous state belief (S_i_) and probabilistic color belief (C_i_), (2) a model with probabilistic state belief (S_i_) and dichotomous color belief (C_i_), and (3) a model in which both state belief (S_i_) and color belief (C_i_) are dichotomous. Here, dichotomous belief refers to the fact that the information is represented in an all-or-none fashion that does not consider uncertainty. The models were evaluated using two criteria. First, we compared their ability to predict the task performed by participants correctly. Second, we compared their BIC scores.

### 2.6 fMRI data acquisition

Imaging data was collected at the Magnetic Resonance Research Facility at the University of Iowa using a 3T GE SIGNA Premier scanner with a 48-channel head coil. Structural images were acquired using a multi-echo MPRAGE sequence (TR = 2348.52ms; TE = 2.968ms; flip angle = 8°; field of view = 256*256; 200 sagittal slices; voxel size = 1mm^3^). Functional images were acquired using an echo-planar sequence sensitive to blood oxygenated level-dependent (BOLD) contrast (multiband factor = 2; TR = 2039ms; TE = 30ms; flip angle = 75°; voxel size = 2mm^3^).

### 2.7 fMRI data preprocessing

All fMRI data were preprocessed using fMRIPrep version 23.2.0 (Esteban et al., 2019) to reduce noise and transform data from subject native space to the ICBM 152 Nonlinear Asymmetrical template version 2009c for group analysis (Fonov et al., 2011). Preprocessing steps include bias field correction, skull-stripping, co-registration between functional and structural images, tissue segmentation, motion correction, and spatial normalization to the standard space. As part of preprocessing, we obtained nuisance regressors including rigid-body motion estimates (three translations and three rotations), cerebral spinal fluid (CSF), and white matter (WM) noise components from the component-based noise correction procedure (Behzadi, Restom, Liau, & Liu, 2007). We also included derivatives and squares of these components. These nuisance regressors were entered in subsequent regression models to reduce the influences from noise and artifacts. We also censored the first three TRs of each functional run and any high motion TRs (framewise displacement greater than 0.5) to remove them from subsequent regression models. We did not perform any spatial smoothing when preprocessing the data.

### 2.8 Statistical analyses

#### 2.8.1 Mixed effects models predicting response times

To examine how trial-by-trial variations in cognitive integration influence behavior, we conducted two separate mixed effects models predicting participant response times (RTs). The models used either (1) the integrated uncertainty (i.e., entropy of the joint distribution) or (2) the uncertainty of the task belief, which was the output of the joint distribution. Both models included random intercepts for subjects. Response times faster than 200 milliseconds and slower than 3500 milliseconds were excluded from analyses. The first trial of each block was also excluded. The entropy of the joint distribution was winsorized to address outliers (values in the bottom 5% and top 95% were replaced with the values at 5^th^ or 95^th^ percentile, respectively). Winsorized entropy and task uncertainty values were approximately normally distributed and homoscedastic. We performed two separate mixed effects regressions due to the high correlation between entropy and task uncertainty (r = .76). We predicted that both greater entropy and greater task uncertainty would predict slower RTs.

We conducted further control analyses with additional behavioral mixed-effects regression analyses of RTs. We compared the above regression using integrated uncertainty (entropy) to predict RTs to a regression using the color uncertainty to predict RTs and a regression using both integrated uncertainty (entropy) and color uncertainty to predict RTs. BIC scores were used to compare models because they are the most conservative when it comes to model overfitting and therefore best suited to determine whether entropy alone is truly the best fit for the data. We predicted that integrated uncertainty (entropy), rather than color uncertainty, would provide the strongest explanatory power for RTs and that including color uncertainty would not provide a significant improvement in explanatory power that justifies the inclusion of an additional predictor (i.e., worth the risk of overfitting).

#### 2.8.2 fMRI Parametric modulation analyses

To relate our model parameters to fMRI data, we conducted parametric modulation analyses to identify brain regions where BOLD response amplitudes vary systematically with key parameters from our computational model. These parameters included (1) the uncertainty of the model predicted inputs-to-integration (state belief and color belief) and the integrated uncertainty (entropy) and (2) prediction errors of inputs (state and color beliefs) and outputs (task belief) and the state derivative, which measured the magnitude of the change in state belief across trials. Details on how these parameters were calculated can be found in section 2.4. These analyses aimed to capture the representational structure, including its uncertainty estimates, and computational processes underlying cognitive integration.

We ran three different mass univariate GLMs using AFNI’s 3dDeconvolve with a voxel-wise restricted maximum likelihood (REML) estimate of an ARMA (1,1) temporal correlation model using AFNI’s 3dREMLfit (Cox, 1996). The first GLM included z-scored state uncertainty, color uncertainty, and entropy as parametric regressors. The second GLM included z-scored task, state, and color prediction errors as parametric regressors. The third GLM included state derivative as a parametric regressor. As mentioned in methods section 2.7, nuisance regressors and censor information were included in all GLMs to reduce the impact of head motion and other noise components on the estimated hemodynamic response function (HRF). These analyses allowed us to assess how each modulator (entropy/uncertainty, prediction error, and state derivative) modulates the HRF magnitude across brain regions on a trial-by-trial basis, beyond trial-averaged effects from the event epochs.

Finally, we conducted mass univariate t-tests on the resulting beta values. These tests identified structures where changes in entropy/uncertainty, prediction error, or state derivative estimates corresponded to significant changes in the BOLD response. To correct for multiple comparisons, we used 3dClustSim with the residuals from the REML estimate to estimate the minimum cluster size for a corrected cluster-level threshold. Significant clusters for parametric modulation are based on these calculated minimum cluster sizes of 128 voxels (cluster p-value threshold of 0.005 and a voxel-level p-value threshold of .05).

#### 2.8.3 fMRI probabilistic decoding analyses

We further conducted decoding analyses to identify the neural substrates encoding (1) the input sources of information, state belief (S_i_) and color belief (C_i_) and (2) the output of this integrated product, task belief (T_i_).

We first obtained trial-by-trial estimates of BOLD response amplitudes using a least-square-sum approach implemented in AFNI’s 3dLSS (Cox, 1996). As mentioned in methods section 2.7, nuisance regressors and censor information were included to reduce the impact of head motion and other noise components on the trial-by-trial BOLD estimates. Decoding was performed via linear regression (Python package scikit-learn; (Pedregosa, 2011)), with trial-level model estimated participant beliefs (state belief, S_i_, color belief, C_i_, or task belief, T_i_) serving as the predicted variable, and the trial-level, voxel-wise beta estimates from 3dLSS as features. This decoding analysis was conducted separately for three distinct 3dLSS analyses, each time-locked to a different task phase (cue onset, probe onset, and feedback onset). The decoding results in section 3.2.1 focus on the cue onset because integration should occur following cue presentation. Whereas results in section 3.2.2 focus on the cue and probe periods because this section addresses representations related to both integration and task execution.

Participant beliefs ranged from zero to one, where boundary values (i.e., near 0 or 1) indicated higher confidence, i.e., lower uncertainty, and values near 0.5 reflected lower confidence, i.e., greater uncertainty (Fig. 3B). To better meet the assumption of normality, these beliefs were log-transformed prior to decoding. We applied a whole-brain searchlight decoding approach with an 8mm radius for each searchlight sphere. Within each searchlight sphere, voxel features were standardized by removing the mean and scaling to unit variance. For decoding, trial-wise betas were split into five groups based on the five functional runs, and a leave-one-group-out cross-validation approach was used to predict the trial-wise beliefs. To assess decoding performance, we correlated the predicted trial-level belief values (decoding output) with the actual model estimated belief values (decoding input), identifying brain regions where the predicted values closely matched the true values (i.e., where decoding performance was higher). Finally, we conducted mass univariate t-tests that tested if these correlation values across subjects were significantly different from zero. To correct for multiple comparisons, we used 3dClustSim with the residuals from a voxel-wise restricted maximum likelihood (REML) estimate of an ARMA (1,1) temporal correlation to estimate the minimum cluster size. Significant clusters for decoding results are based on these calculated minimum cluster sizes of 128 voxels (cluster p-value threshold of 0.005 and a voxel p-value threshold of .05).

#### 2.8.4 Joint probability decoding

Similar to probabilistic decoding (section 2.8.3), we first obtained trial-by-trial estimates of BOLD response amplitudes using a least-square-all approach implemented in AFNI’s 3dLSS (Cox, 1996). We then conducted a multinomial logistic regression (Python package scikit-learn; (Pedregosa, 2011)), with the four possible combinations of state and color belief serving as the four categorical predicted variables, and the beta estimates from 3dLSS as features. As mentioned in methods section 2.7, nuisance regressors and censor information were included to reduce the impact of head motion and other noise components on the trial-by-trial BOLD estimates. This analysis was also conducted separately for the distinct 3dLSS analyses, each time-locked to a different task phase (cue onset and probe onset).

We applied a whole-brain searchlight decoding approach using the same feature extraction and cross-validation settings described in section 2.8.3. The trained multinomial logistic regression model predicted possible classes: state1–color1, state1–color2, state2–color1, and state2–color2. Prediction accuracy was tested against the chance-level decoding performance of 0.25 using mass-univariate t-tests across subjects. Multiple-comparisons correction was performed 3dClustSim with the residuals from a voxel-wise REML estimate of an ARMA (1,1) temporal correlation to estimate the minimum cluster size. Significant clusters for decoding results are based on these calculated minimum cluster sizes of 128 voxels (cluster p-value threshold of 0.005 and a voxel p-value threshold of .05).

#### 2.8.5 Functional network hub analysis

To assess the connector hub-like properties of regions fluctuating with entropy/uncertainty, we calculated participation coefficient (PC) values, which measure how diversely connected a node is within a network (Gratton, Nomura, Perez, & D’Esposito, 2012; Hwang, Shine, Bruss, Tranel, & Boes, 2021). Note, because resting state was not collected with the participant sample reported here, we ran these network hub analyses on an independent validation resting state dataset, previously reported in (Holmes et al., 2015; Hwang, Bertolero, Liu, & D’Esposito, 2017; Reber et al., 2021).

Participation coefficients were calculated on a voxel-wise basis using existing resting state fMRI data (Reber et al., 2021). By leveraging this intrinsic functional connectivity dataset, we can directly compare the spatial distribution of connector hub regions identified using intrinsic, task-free connectivity, to the task-related, entropy-modulated evoked response patterns reported in the current study. This result is implemented using data and methods developed in Reber et al. (2021), and briefly described here.

The PC value for each ROI, PC*_i_* is defined as

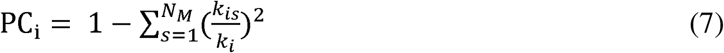

where *k_i_*is the sum of the total functional connectivity weight for ROI *i*, *k_is_*is the sum of the functional connectivity weight between ROI *i* and the cortical network *s*, and *N_M_* is the total number of cortical networks.

These PC values ranged from 0 to 1, with a greater value indicating grater connectivity (i.e., greater “hub”-like properties). Plots were thresholded with a voxel-wise value of 0.65 in order to visualize regions with connector hub-like properties.

### Data availability

Code and data are available at https://github.com/HwangLabNeuroCogDynamics/FPN-Hubs-Integrated-Uncertainty

## Results

To address cognitive integration of goal information, we designed a task paradigm in which participants integrated one external and one internal source of stochastic information to determine the correct task for each trial (Fig. **2**). The external source was a perceptual array of red and yellow dots (Fig. **2C**), while the internal source was a hidden state that covertly changed every 10 to 20 trials (Fig. **2A**). Participants performed either the “Face task”, a gender judgment, or “Scene task”, a place judgement (Fig. **2D**). Participants had to rely on feedback to infer whether the hidden state had changed (Fig. **2E**) because there was no other external indication of state transitions. This paradigm was utilized in conjunction with a computational model, which was designed to capture non-observable aspects of cognitive integration, and fMRI to investigate the neural underpinnings of cognitive integration.

### 3.1 The integration of probabilistic representations into a joint probability distribution drives response selection

We developed a Bayesian model (Fig. **3A**) to capture the computational processes underlying cognitive integration in this paradigm. This model estimates two probabilistic distributions, the volatile hidden state and the ambiguous perceptual cue, which are integrated into a joint probability distribution. This model was applied separately to each subject’s behavioral data. By varying the “noisiness” (i.e., ambiguity/volatility) of these inputs across trials, we manipulated integration uncertainty (i.e., entropy). The joint probability provided a direct measure of integration while the entropy of the joint distribution offers an indirect measure of overall, or integrated, uncertainty. This integrated uncertainty (entropy) in turn influenced the model output (i.e., task to be performed). Furthermore, by tracking uncertainty in both inputs and outputs, the model allowed us to investigate how these representations are updated on a trial-by-trial basis using prediction errors and Bayes’ rule. In short, the model outlines a clear framework that generates trial-by-trial predictors for input representations, their integrated product (i.e., joint probability distribution), the integrated uncertainty (entropy), and the resulting task output representation.

The key feature of our proposed model is that both sources of stochastic information are encoded probabilistically, enabling not only their categorical beliefs, but also their uncertainty to be integrated into a joint representation that predicts task output. To test this probabilistic integration assumption, we compared our model to four alternative models that lacked this feature: (1) dichotomous state and probabilistic color (dSpP), (2) probabilistic state and dichotomous color (pSdP), (3) both state and color dichotomous (dSdP), and (4) a model identical to the. pSdP model but with random guessing under highly ambiguous color conditions (pSjP). This pSjP model was included to capture the possibility that the participant used two different strategies (i.e., random guessing for high color uncertainty and categorical judgments at lower uncertainty).

In these alternative models, representing state or color as dichotomous means beliefs do not include uncertainty, only a categorical decision. Integration in these models thus occurs as a sequential decision process without forming a full probabilistic joint representation. Consequently, belief updating is also treated as an all-or-none process in which beliefs are either entirely revised or remain unchanged. In contrast, the proposed model accounts for graded uncertainty in both state and color beliefs. If the proposed model, where both sources are probabilistic, outperforms the alternative models, this would support the use of entropy in the joint distribution as a measure of integration uncertainty in behavioral and neuroimaging analyses.

We fit all models to behavioral data from the 38 usable subjects who completed the fMRI task. Our proposed probabilistic model (pSpP) showed the lowest prediction error rate (23.08%) compared to alternative models (pSdP: 25.03%, dSpP: 33.70%, dSdP: 24.93%, pSjP: 50.04%; Fig. **4A**). Bayesian information criterion further showed strong preference of the proposed integration model (BIC; Fig. **4B**; protected exceedance probability = 0.945, Bayes Omnibus risk < 0.001). Notably, the alternative model assuming a mixture of random guessing and dichotomous color (pSjP) performed the worst, while the other two models assuming dichotomous color (pSdP and dSdP) showed the closest performance to the proposed model. This likely reflects that color perception is nearly, but not completely, categorical, as indicated by the steep slopes of the sigmoid function used to model color. Overall, these results provide empirical support for the key assumptions of our integrative model.

**Fig 4.**
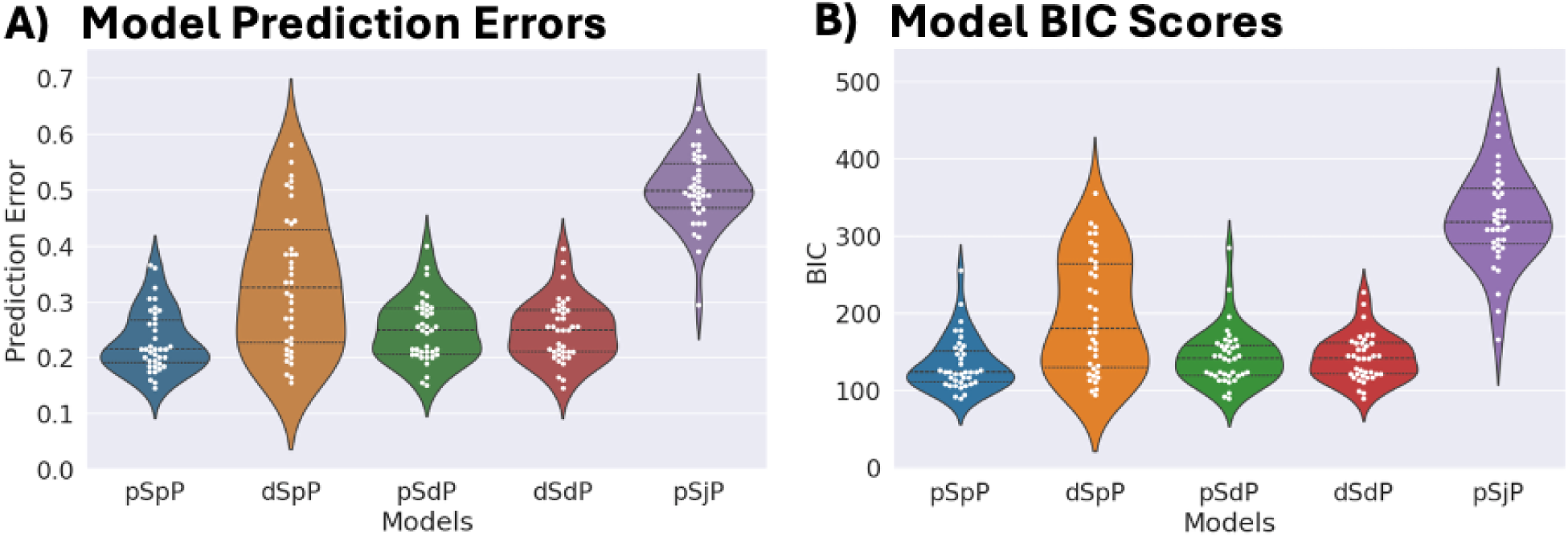
Integration Uncertainty and RTs. (**A**) Prediction error (proportion of trials) for each computational model. Each dot represents an individual subject’s prediction accuracy. The probabilistic model (pSpP) had the lowest prediction error of all the models. (**B**) BIC scores for each computational model. The probabilistic model (pSpP) had the lowest BIC score. Each dot represents an individual subject’s BIC score.

To further test the behavioral relevance of our model, we performed additional analyses on reaction times (RTs). If entropy of the joint distribution indexes integrated uncertainty, greater entropy should predict slower RTs. To test this, we performed a mixed-effects regression with RT as the dependent variable, entropy as a fixed effect, and participants as random intercepts. Higher entropy significantly predicted slower RTs (β = 0.028, t(7572)=3.064, p=.002, Cohen’s d=0.50). Since task belief is derived from this joint distribution, greater task uncertainty should similarly predict slower RTs. A second mixed-effects regression confirmed that task uncertainty significantly predicted slower RTs (β = 0.038, t(7547)=2.257, p=.024, Cohen’s d=0.37). Together, these findings support that integrated uncertainty (entropy) and task uncertainty significantly influence RTs, thus supporting the behavioral relevance of our computational model.

We further tested whether input-level uncertainty, specifically color uncertainty, explained additional variance in RTs beyond entropy. When both entropy and color uncertainty were entered into the same mixed-effects model, entropy remained a robust predictor (β = 0.049, SE = 0.013, z = 3.74, p < .001), whereas color uncertainty was not significant (β = 0.074, SE = 0.050, z = 1.48, p = .14, Cohen’s d ≈ 0.02). Model comparison confirmed that the entropy-only model provided the best fit (BIC = 6259.43), outperforming the color-uncertainty model (BIC = 6279.33) as well as the combined model including both predictors (BIC = 6294.14). Together, these behavioral analyses indicate that integrated uncertainty (entropy) and task uncertainty significantly influence RTs, whereas color uncertainty alone provides no additional explanatory power once entropy is considered.

### 3.2 Stochastic inputs encoded in distinct and distributed regions while integrated uncertainty (entropy) converges in frontal and parietal hub regions

Our model-driven approach enabled us to investigate the neural substrates associated with each component of integration process—inputs, joint representation, output, prediction errors, and updating. Our framework addresses three critical aspects: (1) how distinct information sources are integrated into a unified joint representation, (2) how integration uncertainty, quantified as entropy, biases the resulting task output, and (3) how trial-level feedback drives the updating of input representations. Guided by our model, we ran a series of whole brain multivariate fMRI analyses to investigate every stage of integration. First, we decoded trial by trial beliefs for input beliefs, state and color, and the output task belief to identify the neural representations of input and output information. Next, we used model uncertainty estimates to identify regions where activity scaled with input or integrated uncertainty. We then decoded the joint state–color probability and added its entropy as a parametric regressor to reveal neural representations of the integrated product. To investigate updating processes, we identified structures whose activity varied with prediction errors and state level belief changes. Together, these analyses aimed to reveal the complete cognitive integration process.

#### 3.2.1 Distinct regions encode non-observable and observable inputs

The inputs for integration were an (1) externally presented color cue and (2) internally maintained and continuously tracked state. This continuously tracked state determines the current mappings between color (red or yellow) and task-to-perform (face or scene tasks). The state is continuously updated based on trial-level feedback because no external signal directly indicates its status, while the color cue changed trial-by-trial. These differences led us to predict dissociable neural substrates for encoding each input.

Prior human neuroimaging research suggests the orbitofrontal cortex (OFC) (Moneta, Grossman, & Schuck, 2024; Nassar, McGuire, Ritz, & Kable, 2019; Schuck, Cai, Wilson, & Niv, 2016) and the mediodorsal thalamus (Chen, Leach, Hollis, Cellier, & Hwang, 2024) encodes state like information. We hypothesized that OFC and medial thalamic activity would correlate with model-derived trial-level state beliefs (see Fig. **5A** or methods section 2.8.3 for decoding details). Following cue onset, we found significant clusters in the left primary visual cortex, right middle frontal gyrus (MFG), left primary sensory cortex, right inferior parietal lobule (IPL), and, as expected, the mediodorsal thalamus (Fig. **5B**).

**Fig 5.**
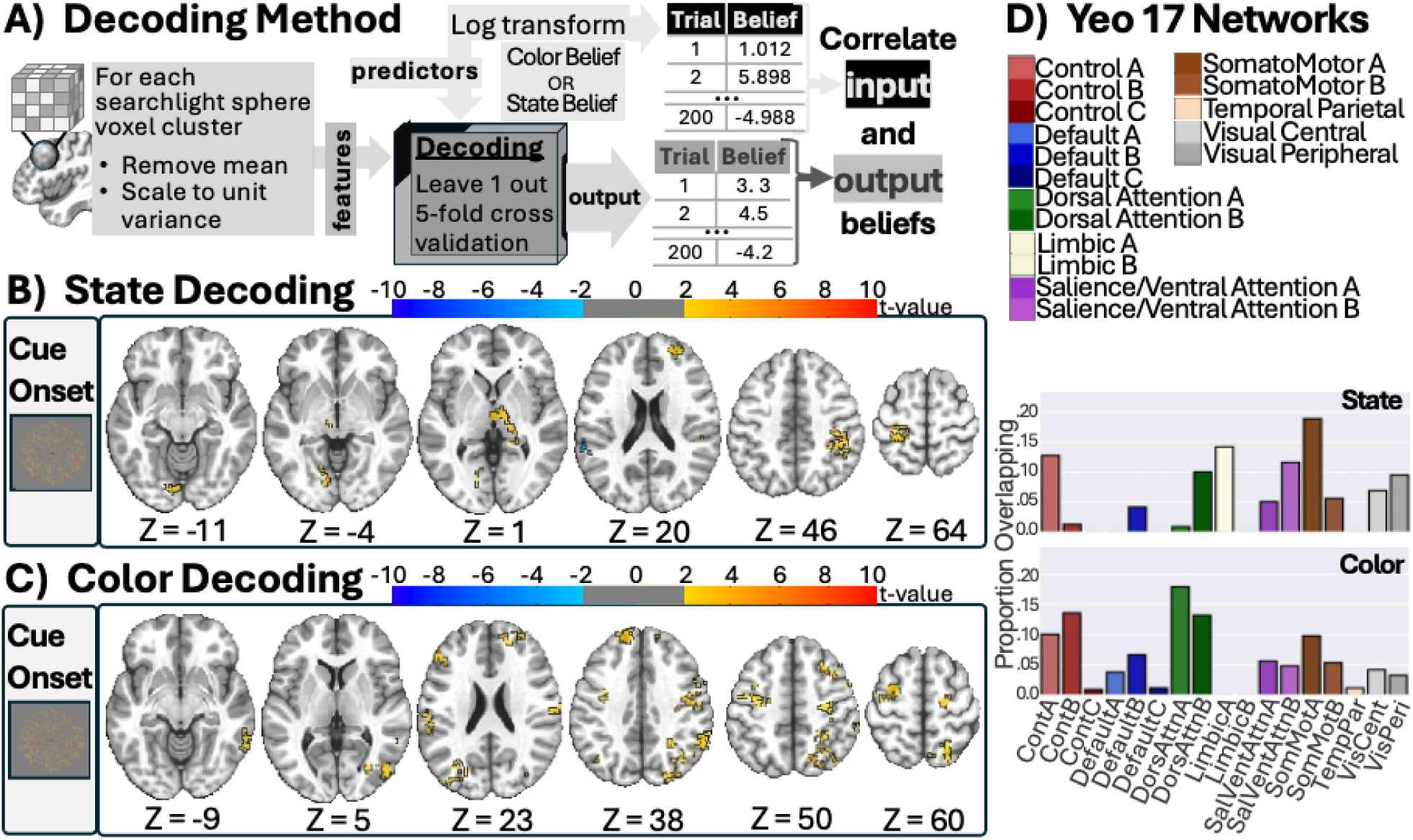
Input Decoding. (**A**) Depiction of decoding method for color and state belief. Voxel clusters were generated with a searchlight method and decoding was performed with a leave one out cross validation approach. Decoding results for state (**B**) and color (**C**) belief during cue epochs showing distributed clusters throughout frontal and parietal as well as some lateral or medial clusters in temporal and occipital regions. Results based on minimum cluster size of 128 voxels (cluster p-value threshold of 0.005 and a voxel-level p-value threshold of .05). (**B**) Yeo 17 functional networks color key and bar plots in depict the proportion of voxels (only from the significant clusters) that overlap with each of the 17 functional networks. The Schaefer 400 ROI atlas was used to obtain the 17 functional networks.

Previous studies showed that externally presented cues, linked to task rules, elicited activity in the several lateral frontal and intraparietal regions, some overlapped with the frontoparietal and dorsal attention networks (Cole et al., 2013; Corbetta et al., 1998). Given that the color belief was derived from externally presented perceptual information (the cue at the start of the task trial) and is linked to task rules (cue color informs the subsequent task-to-perform), we predicted that trial-level color beliefs would be decodable from the lateral occipito-temporal, intraparietal sulcus (IPS), and lateral PFC. Consistent with this prediction, decoding analyses following cue onset revealed distributed clusters across prefrontal, temporal, limbic, occipital, and parietal cortices (Fig. **5C**), including significant clusters in right lateral occipito-temporal and bi-lateral frontal and parietal regions: bilateral superior frontal gyrus (SFG), right MFG, left inferior frontal gyrus (IFG), superior parietal lobule (SPL), and bi-lateral IPS.

To determine which functional networks were involved in encoding these representations, we quantified the overlap between decoded voxels and the Schaefer–Yeo 17 network parcellations (Schaefer et al., 2018). Regions where state information was decoded overlapped primarily with the limbic, somatomotor, frontoparietal, and salience/ventral attention networks (Fig. **5D****-top**). In contrast, regions where color information was decoded showed highest overlapped with the frontoparietal control and dorsal attention networks (Fig. **5D****-bottom**). This indicates that perceptual beliefs are not confined to visual cortices but extend into higher-order control and attention systems, consistent with our manipulation that color belief involves resolving ambiguous color presentation. Although state and color engaged some overlapping networks, their dominant associations were distinct. The observed somatomotor overlap for both could be related to the fact that task identity was linked to opposite hands and that both color and state information are needed to determine the task-to-perform (e.g., in state 1, red cues the face task; in state 2, red cues the scene task).

Finally, to statistically compare the strength of state and color information encoded in distinct regions, we compared state and color decoding performance in independently defined regions of interest (Schaefer et al., 2018; Yeo et al., 2011). A set of ROIs showed significant differences (p <0.05, uncorrected) between state and color decoding (Supplementary Figure S1). Specifically, the right precentral sulcus, the right inferior parietal lobule, the right IPS, and the left rostral and caudal middle frontal gyrus showed stronger color decoding. The medial thalamus showed stronger state decoding when compared to color. This pattern of dissociation raises a critical question: how does the brain integrate these disparate representations to form a unified task output?

#### 3.2.2 Prefrontal and parietal regions encode the integrated joint representation, its associated uncertainty, and the output task belief

In our framework of cognitive integration, the joint representation is defined as the posterior distribution formed by integrating multiple probabilistic inputs. This joint probability, which reflects specific combination of color and state, can be used to predict the output of integration, expressed as a task belief that determines which task to perform. Following cue onset, we expected preparatory activation in brain regions encoding task belief. We hypothesized that the joint probability and task belief would be decoded from frontal and parietal regions associated with cognitive control, such as more caudal regions of MFG and IPS.

##### 3.2.2.1 Joint Probability and Task Belief are encoded in overlapping brain regions

In line with our predictions, decoding analyses of joint probability identified significant clusters in MFG and IPS, as well as parahippocampal (with cue onset) and hippocampal (with probe onset) regions and FFA/PPA (Fig. 6A). Robust clusters were also observed in the supplementary motor cortex regions across the primary motor and sensory cortices towards IPS. Patterns were highly similar across cue and the probe epochs. Furthermore, for decoding task belief, distributed clusters were found in regions where the joint probability was decoded (Fig. 6B). Of particular note, task belief was decoded from the same regions spanning from supplementary motor cortex regions across the primary motor and sensory cortices into the IPS. This pattern may reflect the fact that participants could preparatorily activate the left or right hand depending on which task they believed they should perform.

**Fig 6.**
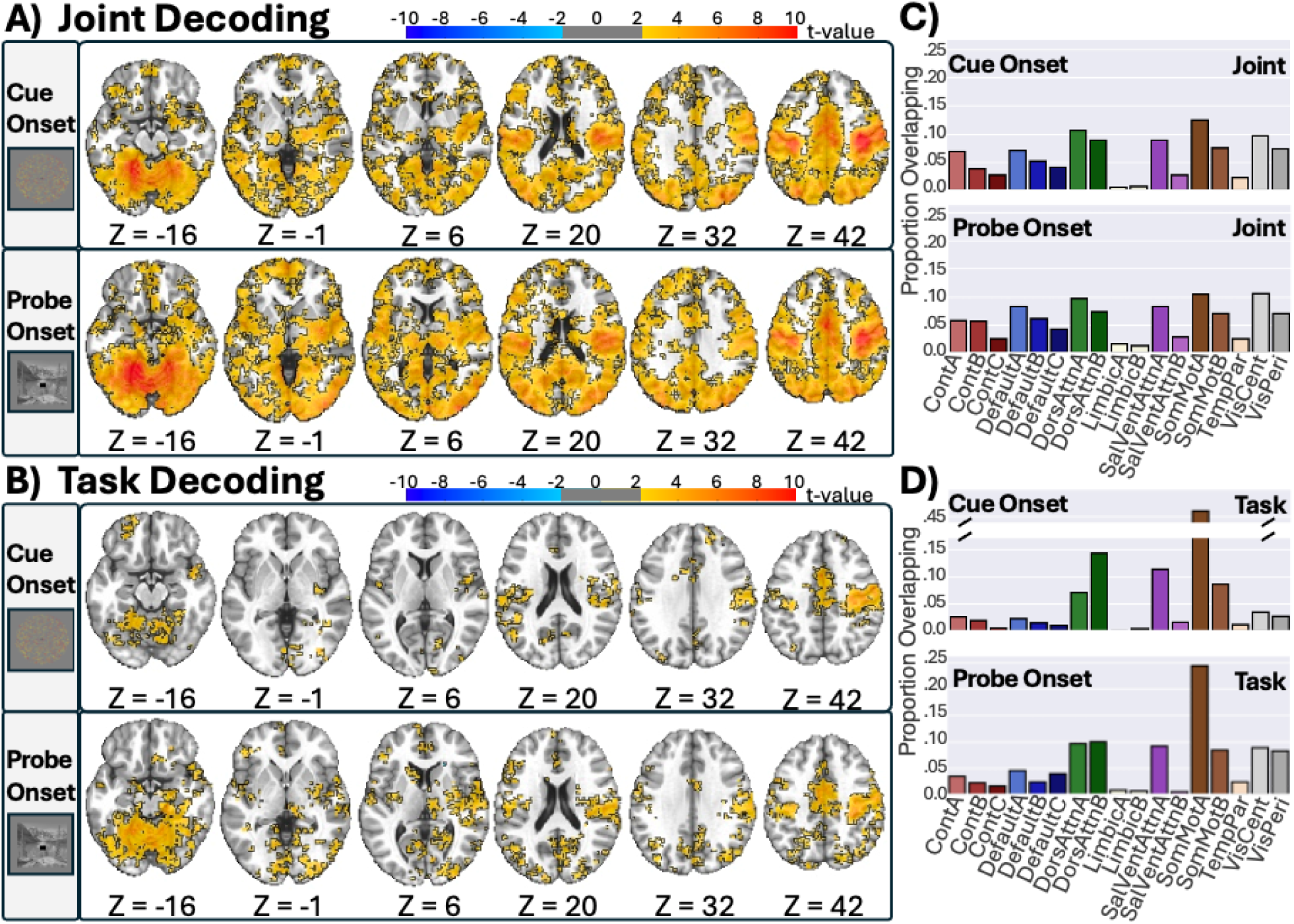
Integration and Output Decoding. Decoding results (**A-B**) for joint probability (**A**) and task belief (**B**) during cue and probe epochs based on a minimum cluster size of 128 voxels (cluster p-value threshold of 0.005 and a voxel-level p-value threshold of .05). Decoding for task belief was performed with the same method depicted in Fig. 5A. Decoding for the joint probability distribution was performed similarly to task belief, but with multinomial logistic regression model predicting the specific color and state combination. Bar plots in **C-D** depict the proportion of voxels (only from the significant clusters) that overlap with each of the 17 functional networks. The Schaefer 400 ROI atlas was used to obtain the 17 functional networks. The face stimuli in the probe example image is of an author on this manuscript.

We repeated the same network overlap analyses to determine which networks encode the joint probability information and the task output. Network overlap analyses revealed that regions with decodable joint probability information were distributed across nearly all 17 networks (Fig. **6C**), with the limbic network showing the least involvement. Network engagement remained largely consistent across cue and probe epochs. Regions from which task belief was decoded were primarily located in the somatosensory/motor networks (Fig. **6D**).

##### 3.2.2.2 Prefrontal and parietal regions responded to the integrated uncertainty (entropy of the joint representation)

We further conducted a parametric modulation analysis to identify regions where activity scales with input (state and color) uncertainty and integrated uncertainty (entropy). Entropy captures the combined uncertainty of state and color inputs; entropy remains low when both are certain, high when both are uncertain, and intermediate otherwise. Following cue onset, regions including MFG, mPFC, precentral sulcus, IPS, and SFG exhibited significant modulation with integrated uncertainty, or entropy of the joint distribution (Fig. **7A**). When including all three uncertainty estimates in the same model, there were no significant clusters for input uncertainty, only for integrated uncertainty (entropy). In short, this suggests that the brain encodes an overall integrated uncertainty from multiple inputs. Finally, network analyses (Fig. **7C**) revealed that regions fluctuating with integrated uncertainty (entropy) primarily overlapped with the frontoparietal control, dorsal attention, and salience/ventral attention networks.

**Fig 7.**
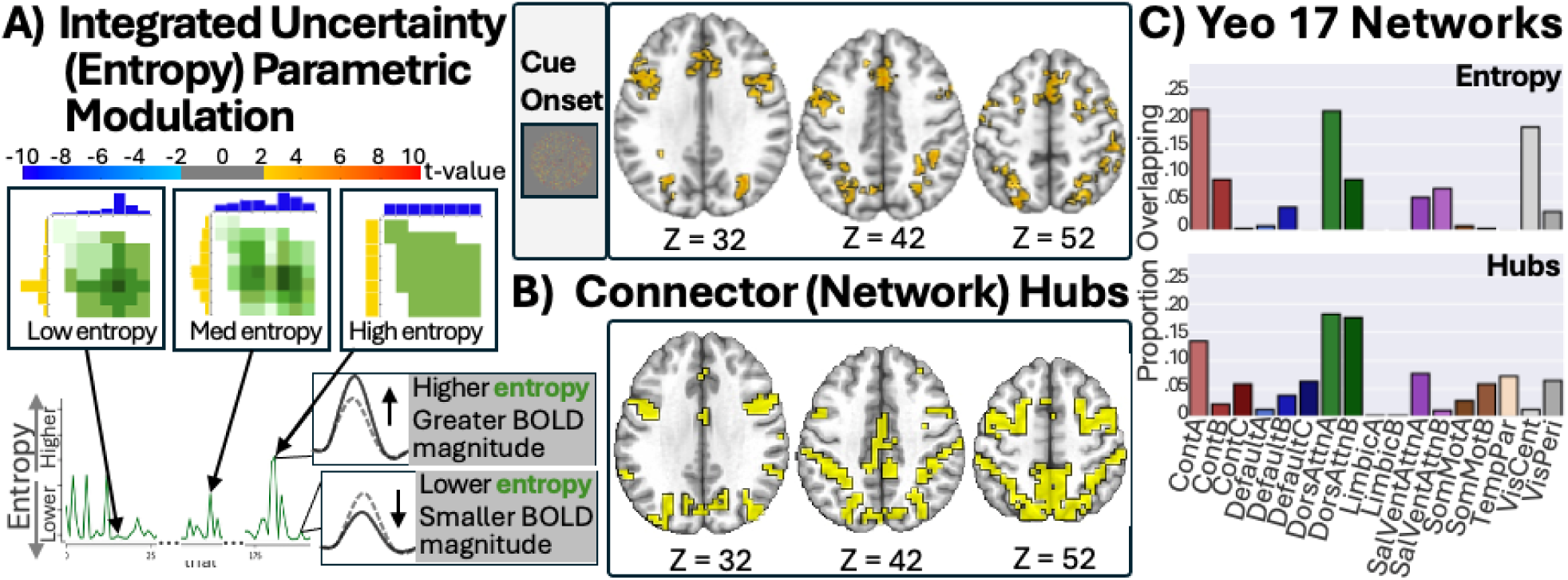
Integrated Uncertainty (Entropy) and Network Hubs. (**A**) Parametric modulation analyses with z-scored input (state and color) uncertainty and integrated uncertainty (entropy) revealed robust significant clusters for integrated uncertainty (entropy) based on a minimum cluster size of 128 voxels (cluster p-value threshold of 0.005 and a voxel-level p-value threshold of .05). (**B**) Network hub calculations show that many of the same frontal and parietal regions that encoded integrated uncertainty (entropy) are also hub regions, i.e., highly connected to many networks. Network hubs defined as PC value > 0.65. (**C**) Bar plots depict the proportion of voxels (only from the significant clusters) that overlap with each of the 17 functional networks. The Schaefer 400 ROI atlas was used to obtain the 17 functional networks.

##### 3.2.2.2 Network hub regions encode integrated uncertainty (entropy of the joint representation)

Our integration framework predicts that regions that integrate observable and non-observable information should exhibit network hub-like properties. These hubs regions are expected to track the integrated uncertainty of multiple task-relevant inputs from distinct information channels. Achieving this would require broad distributive connectivity with multiple networks. To test this prediction, we computed voxel-wise participation coefficient (PC) scores using a previously developed method (Reber et al., 2021). PC is a widely used metric do identify network hubs in the human brain (Gratton et al., 2018; Hwang et al., 2021; Power et al., 2013), its value ranged from 0-1 and we a priori defined hubs as having a PC > 0.65. Based on these criteria, MFG and IPS are network hub regions (Fig. 7B). Moreover, there is a strong overlap between regions fluctuating with integrated uncertainty (entropy) and network hub regions (Fig. **7C**). Finally, there was strong overlap between regions that fluctuated with integrated uncertainty (entropy) and hub regions in the control and dorsal attention networks. These findings further support our account in which network hub regions, specifically, leverage integrated uncertainty (entropy) from diverse information sources to bias task output.

In sum, frontoparietal hub regions, such as MFG, mPFC, precentral sulcus and IPS, have thus far shown two key findings. First, their activity fluctuated with a measure of integrated uncertainty (entropy of the joint distribution). Second, these regions showed strong decoding of the joint probability (representing the integrated inputs), and the task belief (output). One interpretation based on these observations is that these regions leverage integrated uncertainty (entropy) from task-relevant input information sources and then utilize this signal to bias the output from integration.

#### 3.2.3 Integrated uncertainty (entropy) drive prediction errors and updating of latent state belief

Previous sections have focused on how the integration of information predicts task output. However, our account of integration would be incomplete without understanding how input representations are updated following feedback. In our model, the state belief requires continuous monitoring across trials and updating because no direct external signal indicates state. Updating state belief thus involves evaluating: (1) the relationship between the received feedback and the performed task, (2) the joint representation of state and color beliefs that led to the task decision, and (3) the current state belief. Thus, state updating required multiple information sources to generate prediction errors and update state belief.

To assess updating processes, we first investigated brain regions where activity was modulated by both output (task) and input (color and state) prediction errors following feedback (Fig. **8A**). Presumably, incorrect feedback first triggers a task prediction error, which will then trigger a re-evaluation of the integrated product and the inputs to integration (color and state belief). Additionally, we examined regions where activity varied with the trial-to-trial change in state belief, termed the state derivative (Fig. **8D**). These state derivative analyses were time-locked to the cue onset on trial N to investigate updating processes occurring following feedback on the previous trial (trial N-1).

**Fig 8.**
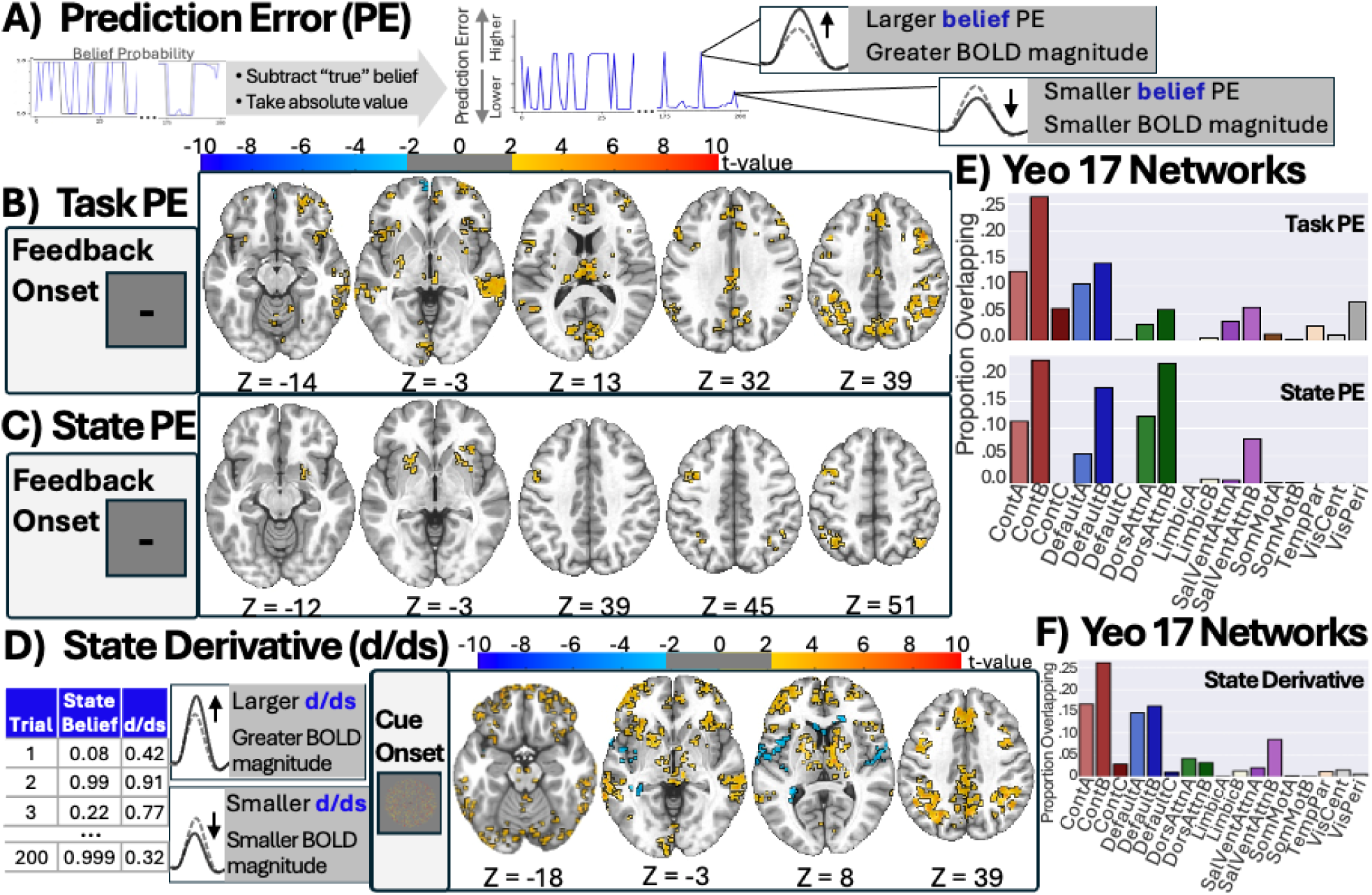
Prediction Errors and Updating Parametric Modulation. (**A**) Depiction of parametric modulation analyses with (**B**) task prediction errors and (**C**) state prediction errors at feedback onset. Prediction error calculated by subtracting the “true” task from the model estimated task and taking the absolute values. (**D**) Parametric modulation with state derivative at cue onset, which was calculated by subtracting the state belief on trial N-1 from the state belief on trial N and taking the absolute value. All results based on minimum cluster size of 128 voxels (cluster p-value threshold of 0.005 and a voxel-level p-value threshold of .05). Bar plots (**E-F**) represent the proportion of voxels (significant clusters only) overlapping with each of the 17 functional networks. The Schaefer 400 ROI atlas was used to obtain the 17 functional networks.

##### 3.2.4.1 Greater belief uncertainty associated with greater prediction errors in prediction error circuits

We focused on z-scored input and output prediction errors while controlling for general error processing (i.e., whether feedback was correct or incorrect). Using feedback-locked data, we predicted that mPFC voxels would show increased BOLD responses with larger prediction errors, consistent with its known role in conflict and error monitoring (Botvinick, Braver, Carter, & Cohen, 2001; Freund, Bugg, & Braver, 2021; Jiang, Wagner, & Egner, 2018; Rutledge, Dean, Caplin, & Glimcher, 2010). As expected, task prediction errors showed greater activation in mPFC (Fig. **8B-C**). Color prediction errors did not show robust clusters. Beyond these expected regions, we also found that regions in the posterior parietal cortex (PPC), lateral PFC, and subcortical structures, such as the basal ganglia, showed a greater magnitude in their BOLD response with greater state and task prediction errors.

##### 3.2.4.2 Greater updating of latent state belief associated with stronger activity in inferior and middle frontal gyrus and subcortical structures

We expected greater changes in state belief from trial N-1 to trial N (i.e., a higher state derivative) to be associated with increased activity in the OFC, a region implicated in tracking state information (Moneta et al., 2024; Nassar et al., 2019; Schuck et al., 2016). Furthermore, given that subcortical structures such as the basal ganglia are involved in working memory updating, we anticipated these regions would also exhibit stronger BOLD activity with larger state derivatives, reflecting greater updating demands. In line with these predictions, both the OFC and basal ganglia (caudate nucleus), as well as the medial thalamus, showed increased BOLD magnitude with greater state derivative (Fig. **8D**).

For both prediction errors (Fig. **8E**) and state derivatives (Fig. **8F**), regions fluctuating with prediction errors or the magnitude of change in state belief showed overlap with control, default mode, dorsal attention, and salient/ventral attention networks. Once again, control networks accounted for the greatest proportion of significant voxels. Together, these findings show that greater uncertainty levels lead to larger prediction error responses in prefrontal, parietal, and subcortical regions, while trial-to-trial changes in state belief recruit OFC, basal ganglia, and thalamus. In short, uncertainty provides the basis for generating prediction errors and drives updating of latent state beliefs across distributed cortical–subcortical circuits.

## Discussion

Our primary goal was to determine the neurocognitive mechanisms supporting cognitive integration —specifically, how stochastic beliefs and perceptual information are integrated to guide behavior. We developed a Bayesian model capturing the full transformation from inputs to outputs by specifying: (1) the representational structure of inputs, (2) the integrated representation (joint probability distribution) encoding the integrated uncertainty (entropy) of these inputs, (3) the probabilistic output (task) belief, and (4) feedback-recruited prediction errors and updating mechanisms.

Our first aim was to identify the neural substrates representing the inputs to the integration process. Probabilistic decoding revealed that perceptual inputs and internally maintained beliefs were represented in largely distinct neural substrates. Color belief was represented in lateral occipitotemporal cortex and in frontoparietal regions typically associated with the FPN and DAN (Cole et al., 2013; Corbetta et al., 1998), including the precentral sulcus, the putative frontal eye fields, premotor regions, and the intraparietal sulcus. Importantly, what we decoded was color belief derived from our computational model, inferred from subjects’ responses, rather than the physical stimulus values. This means the signal reflected the accumulation of perceptual information towards forming a color belief in addition to the raw sensory input. From this perspective, the engagement of frontoparietal circuits is theoretically consistent with their suggestive roles in perceptual decision making and evidence accumulation (Bird, Berens, Horner, & Franklin, 2014; de Lafuente & Romo, 2005; Liu & Pleskac, 2011; Mante, Sussillo, Shenoy, & Newsome, 2013; Ploran et al., 2007). In contrast, state belief was primarily represented in the medial thalamus, anterior prefrontal cortex, and superior parietal cortex, overlapping with networks often described as somatomotor, ventral attention, and frontoparietal. The thalamic finding aligns with evidence that the mediodorsal thalamus encodes contextual information that organizes stimulus–response mappings and exerts hierarchical influence over cortical activity (Lam et al., 2025). In our prior work (Chen et al., 2024), thalamic encoding was also observed when context was tied to a sensory cue, and here we extend this to a design where latent state must be inferred from other perceptual inputs. This novel finding further suggests that the human medial thalamus encodes latent (not directly tied to a sensory cue) state-like representations. At the same time, state decoding in parietal and motor-associated regions may indicate that retrieving state information co-activated well-practiced stimulus–response mappings, and in our design, color inputs specified both the task and its response mappings. This may partly account for the engagement of motor-related areas during state decoding, although this interpretation remains speculative. Although we did not find significant OFC encoding of state at our corrected threshold, we found that the OFC showed robust activity in response to shifts in state representation (state derivative), consistent with prior studies (Konishi, McLaren, Engen, & Smallwood, 2015; Nassar et al., 2019; Vatansever, Menon, Manktelow, Sahakian, & Stamatakis, 2015). The fact that distinct regions encoded color and state belief further underscored the importance of cognitive integration.

Our second and third aims were to identify the neural substrates representing the integrated representation and output task belief. Multinomial logistic regression decoding analyses revealed that the specific combination of inputs— captured by a joint probability distribution—accurately predicted the state-color integration and task output. Significant encoding of these representations was found across distributed regions, spanning across all functional networks. Following probe onset, regions where the joint probability was decoded overlapped with regions encoding output task beliefs, spanning sensory-motor regions that likely reflect preparatory motor activation for the anticipated task. A notable set of regions that showed significant encoding of both task output and state-color integration included the MFG, mPFC, precentral sulcus and IPS. These regions not only encode task output and state-color integration, but also robustly responded to integrated uncertainty (entropy of the joint distribution). Furthermore, these regions showed extensive overlap with FPN, and DAN networks, including nodes that are known hub regions connected to multiple large-scale networks (Bertolero, Yeo, Bassett, & D’Esposito, 2018; Cocuzza, Ito, Schultz, Bassett, & Cole, 2020; Cole et al., 2013; Gratton et al., 2018). These results suggest that frontoparietal regions may detect and combine input uncertainty and then utilize the resulting integrated uncertainty to resolve output competition. The integrated uncertainty and integrated representations encoded in FPN hubs may be converted into a biasing signal that modulates task-relevant regions, ultimately driving the output of a single, unified task belief (Egner & Hirsch, 2005; Miller & D’Esposito, 2005).

Our final aim investigated continuous prediction errors and updating processes. Updating processes recruited subcortical structures (basal ganglia, mediodorsal thalamus) and frontal-parietal regions, with notable hub regions (e.g., MFG, mPFC, IPS) responded to both larger prediction errors and greater integrated uncertainty (entropy). Thus, FPN hubs may further utilize integrated uncertainty (entropy) to bias regions involved detecting discrepancies between task beliefs and true values (prediction errors). This could potentially drive updating of internally maintained state beliefs.

Our findings highlight the importance of uncertainty in integration and task control. Sensory research has long demonstrated that neural activity encodes information probabilistically rather than in an all-or-none fashion (Ma, Beck, Latham, & Pouget, 2006; van Bergen, Ma, Pratte, & Jehee, 2015). However, decision making requires a winner-take-all process where only the most probable option is selected (Vul, Hanus, & Kanwisher, 2009). Our framework proposes that probabilistically encoded inputs are integrated into a joint distribution and generate a conditional probability from which the most likely action is selected. Our results suggest that the human brain leverages the probabilistic information encoded in these inputs and processes them within a common set of frontoparietal regions as opposed to resolving uncertainty at the input level and considering only the categorical information during integration. Moreover, although the output task belief drives motor plans in a winner-take-all manner, this belief also encodes uncertainty to facilitate subsequent updating processes.

Our model suggests that human subjects integrated stochastic inputs by maintaining probability distribution and uncertainty over all possible combinations of information inputs. This approach can offer several advantages. First, it can support cognitive flexibility because every possible input combination already exists within the probabilistic representation. This allows the system to switch between tasks more accurately when the behavioral state changes. Indeed, our behavioral modeling results showed that our probabilistic model outperformed other models that do not maintain probability distribution. Second, probabilistic representations may provide a principled uncertainty-driven control signal (entropy) that can be quantified and utilized for adaptive top-down control. High entropy can signal the need for additional top-down resources. In contrast, low entropy allows strong updating in representations, facilitating flexibility. This mechanism enables the cognitive system to adaptively allocate processing resources based on the integrated information, providing a computational solution for flexible cognitive control under uncertainty.

Our results suggest that this entropy-based mechanism appears to be a core function of cognitive integration. In our study, the entropy of the joint distribution serves as a critical signal that detects competition, noise, and variability across multiple task-relevant input sources. This signal enables the selection of the most likely correct action plan and decision (i.e., task) based on input information. Recent work by Li, Sprague, Yoo, Ma, and Curtis (2024) similarly indicates that uncertainty signals may guide targeted prioritization of multiple representations in working memory, resolving competition between representations. Therefore, the maintenance of probabilistic representations across diverse brain regions and the associated integrated uncertainty (entropy) is integral to both the continuous integration and targeted updating of task-relevant information.

A central implication of our findings concerns the concept of hub regions in human functional brain networks. Many frontoparietal areas in the FPN, DAN, and CON have been labeled “integrative” because of their dense connectivity and central positions in network topology (Bertolero et al., 2018; Cocuzza et al., 2020; Cole et al., 2013; Gratton et al., 2018). However, what specifically makes these regions integrative beyond their connectivity profile has remained unclear. Our results provide a mechanistic account, specifically, we demonstrate a novel entropy-based mechanism of integration. Hubs may serve as convergence zones where uncertainty signals from diverse sources are combined, and to be utilized as a control signal to influence processes across systems. This perspective moves beyond describing hubs as structurally well-connected to identifying what computational role they perform.

In summary, our study demonstrates how the frontoparietal system actively integrates internal and external sources of stochastic information to guide adaptive behavior. By defining the representational structure of the inputs, the integrated product (joint distribution), and the output (task belief), along with their associated uncertainty measures, we tracked information integration from initial encoding to action selection. Results revealed that internal and external inputs were represented in mostly distinct brain structures before converging in FPN hub regions, with entropy serving as a mechanism that guides both task output selection and updating processes. This work provides a computational framework for understanding how the human FPN integrate and update task-relevant information for adaptive, goal-directed behavior.

## Acknowledgments

Author contributions are as follows.

Conceptualization: SCL, KH, JJ

Methodology: SCL, KH, JJ

Investigation: SCL, HH, KH, JJ

Visualization: SCL, KH, JJ

Supervision: KH, JJ

Writing—original draft: SCL, KH, JJ

Writing—review & editing: SCL, KH, JJ

Authors declare that they have no competing interest.

All data and code will be made available upon publication.

Research reported here was supported by a National Institutes of Mental Health grant R01MH122613 (KH) and the Iowa Neuroscience Institute. This work was conducted on an MRI instrument funded by 1S10OD025025-01

