## supplement for "Frontoparietal Hubs Leverage Probabilistic Representations and Integrated Uncertainty to Guide Cognitive Flexibility"

Stephanie C. Leach* *et al.*

**This PDF file includes:**

Figure S1

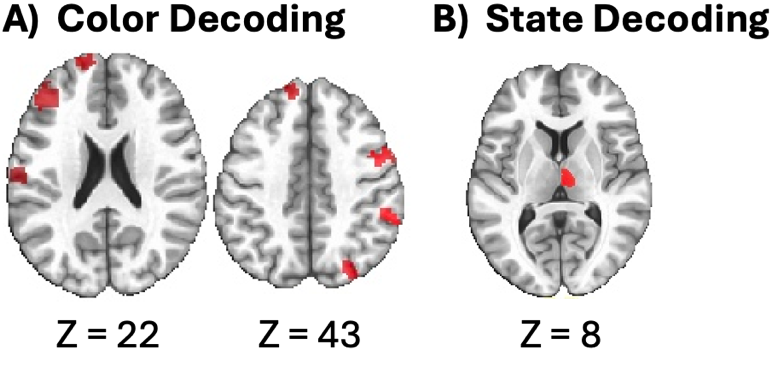

**Figure S1**. **Color vs State Decoding.** Significant ROIs based on peak voxel from (**A**) color decoding or (**B**) state decoding beta maps. To provide an unbiased comparison between state and color decoding, we used an ROI-based cross-analysis procedure that avoided circularity. First, we identified clusters from the group-level color or state decoding maps using cluster-forming thresholds (t ≥ 2.026, minimum cluster size = 128 voxels, NN=2). For each surviving cluster, we located the peak voxel and assigned it to an anatomical ROI using the Morel thalamic atlas (Krauth et al., 2010) and the Schaefer–Yeo 400 cortical parcellations (Schaefer et al., 2018; Yeo et al., 2011). This produced independent sets of ROIs defined from either color or state group-level maps. Next, for each subject, we extracted the mean Fisher-z–transformed decoding accuracy within each ROI, separately for color and state maps. This provided paired values per subject per ROI. We then performed paired-sample t-tests comparing state and color decoding across subjects, within each ROI. The resulting significant ROIs were projected back onto the parcellation masks to generate visualization maps in this supplementary figure (thresholded at t > 2.026, p < 0.05, uncorrected).
